# Effective potential reveals evolutionary trajectories in complex fitness landscapes

**DOI:** 10.1101/869883

**Authors:** Matteo Smerlak

## Abstract

Growing efforts to measure fitness landscapes in molecular and microbial systems are premised on a tight relationship between landscape topography and evolutionary trajectories. This relationship, however, is far from being straightforward: depending on their mutation rate, Darwinian populations can climb the closest fitness peak (survival of the fittest), settle in lower regions with higher mutational robustness (survival of the flattest), or fail to adapt altogether (error catastrophes). These bifurcations highlight that evolution does not necessarily drive populations “from lower peak to higher peak”, as Wright imagined. The problem therefore remains: how exactly does a complex landscape topography constrain evolution, and can we predict where it will go next? Here I introduce a generalization of quasispecies theory which identifies metastable evolutionary states as minima of an effective potential. From this representation I derive a coarse-grained, Markov state model of evolution, which in turn forms a basis for evolutionary predictions across a wide range of mutation rates. Because the effective potential is related to the ground state of a quantum Hamiltonian, my approach could stimulate fruitful interactions between evolutionary dynamics and quantum many-body theory.

**SIGNIFICANCE STATEMENT:** The course of evolution is determined by the relationship between heritable types and their adaptive values, the fitness landscape. Thanks to the explosive development of sequencing technologies, fitness landscapes have now been measured in a diversity of systems from molecules to micro-organisms. How can we turn these data into evolutionary predictions? I show that preferred evolutionary trajectories are revealed when the effects of selection and mutations are blended in a single effective evolutionary force. With this reformulation, the dynamics of selection and mutation becomes Markovian, bringing a wealth of classical visualization and analysis tools to bear on evolutionary dynamics. Among these is a coarse-graining of evolutionary dynamics along its metastable states which greatly reduces the complexity of the prediction problem.

## INTRODUCTION

Darwinian evolution is the motion of populations in the space of all possible heritable types graded by their reproductive value, the fitness landscape [1–3]. In Wright’s vivid words, the interaction of selection and variation enables populations to “continually find their way from lower to higher peaks” [4], thereby providing a universal mechanism for open-ended evolution [5]. Thanks to the explosive development of sequencing technologies, fitness landscapes have now been measured in a variety of real molecular [6], viral [7] or microbial [8] systems. As a result, the goal of *predicting* evolution no longer appears wholly out of reach [8–12]. In essence, if we know the topography of the fitness landscape—its peaks, valleys, ridges, etc.—we should be able to estimate where a population is likely to move next. Making such predictions from high-resolution fitness assays is a central challenge of quantitative evolutionary theory.

An array of measures quantifying the topography of fitness landscapes has been developed in support of this program. Especially important from the dynamical perspective, the *ruggedness* of a landscape can be measured in a variety of ways, some simple but coarse (density of fitness maxima [13], correlation functions [14, 15]), others detailed but more involute (amplitude spectra [16], geometric shapes [17]). As stressed by Kimura [18], *neutrality*—the distribution of plateaus rather than peaks—is another key feature of real landscapes [19] which derives from their high-dimensional nature [20, 21] and can be studied with the tools of percolation theory [22]. The NK(p) [23, 24], Rough Mount Fuji [25, 26], holey [20, 21] and other models in turn provide simple landscapes with tunable ruggedness and/or neutrality, which can be used to fit empirical data and simulate evolutionary trajectories. On these foundations a new subfield of mathematical biology has emerged, the quantitative analysis of fitness landscapes [27].

What these fitness-centric measures fail to capture, however, is the fact that *populations with different mutation rates experience the same fitness landscape differently*. This is already clear if we consider the rate of fitness valley crossings, which strongly depends on the mutation rate [28, 29] and therefore cannot be computed from topographic data alone. But Eigen’s quasispecies theory showed that varying mutation rates can also have a *qualitative* effect on evolutionary trajectories, potentially leading to error catastrophes and the loss of adaptation [30]. More subtly, mutational robustness has been shown to evolve neutrally [31] and to sometimes outweigh reproductive rate as a determinant of evolutionary success (“survival of the flattest”) [32, 33]. These evolutionary bifurcations are not mere theoretical curiosities: lethal mutagenesis—an effort to push a population beyond its error threshold—is a promising therapeutic strategy against certain viral pathogens [34, 35] and perhaps cancer [36].

More fundamentally, these bifurcations imply that, unless mutations are so rare that populations are genetically homogeneous at all times and selection is periodic (the so-called strong-selection weak-mutation (SSWM) limit [37]), evolving populations are not necessarily attracted to fitness peaks. The SSWM assumptions are typically violated in large microbial populations, where multiple mutants often compete for fixation in a process known as clonal interference [38, 39]. In yet stronger mutation regimes, e.g. in RNA viruses, natural selection acts on clouds of genetically related mutants rather than on individuals, and evolution is the intermittent succession of stabilization-destabilization transitions for these clouds [40]. In the presence of neutrality, epochal or “punctuated” evolution can also arise through the succession of slow intra-network neutral diffusion and fast inter-network sweeps [41].

These results raise fundamental questions regarding the *dynamical* analysis of fitness landscapes: When is flatter better than fitter? Where are the evolutionary attractors in a given landscape with ruggedness and/or neutrality? What quantity do evolving populations optimize? Can we estimate the time scale before another attractor is visited? More simply, can we predict the future trajectory of an evolving population from its current location, the topography of its landscape, and the mutation rate?

In this paper I outline a mathematical framework to address these questions in large, asexual populations, for both genotypic (discrete, high-dimensional) and phenotypic (continuous, low-dimensional) fitness landscapes. Inspired by certain analogies with the physics of disordered systems, I show that the selection-mutation process can be understood as a random walk or diffusion in an effective potential—the same kind of dynamics as, say, protein folding kinetics [42]. This representation reduces the *a priori* difficult problem of identifying evolutionary attractors and dominant trajectories in a complex fitness landscape to the much more familiar problem of Markovian metastability [43]. In contrast with another classical Markovian model of evolution, Gillespie’s adaptive walk model [13, 37], my approach is not restricted to the SSWM regime and fully accommodates genotypic and/or phenotypic heterogeneity in evolving populations [38, 39]. Moreover, because the effective potential integrates fitness and mutational robustness in a single function on the space of types, it is also more suited to analyze—and eventually predict—the dynamics of a population than the bare fitness landscape from which it derives.

## RESULTS

### Selection-mutation dynamics

Consider a fitness landscape Φ = (*X*, Δ, *ϕ*), consisting of a space of types *X*, a mutation operator Δ on *X* and a (Malthusian) fitness function *ϕ* : *X* → ℝ. The nature of the landscape is left unspecified: Φ could be a be genotypic landscape, in which case *X* will be a finite graph (usually a hypercube or some more general Hamming graph), and Δ its Laplacian matrix; or Φ could be a quantitative phenotypic landscapes, and then *X* will be a domain of ℝ^*d*^ and Δ a differential operator thereon, usually the Laplacian (if mutational effects are sufficiently small and frequent).

We assume a large asexual population evolving on this landscape according to the continuous-time Crow-Kimura [44] selection-mutation equation

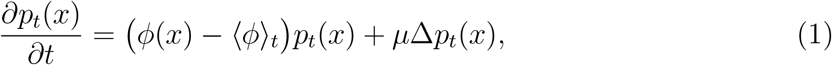

where *p*_*t*_(*x*) is the distribution of types *x* ∈ *X* at time *t*, ⟨*ϕ*⟩_*t*_ the population mean fitness, and *μ* the mutation rate per individual per unit time. In contrast with previous analytical works which focused on finding exact solutions to (1) [45], our goal is to understand the motion of the distribution *p*_*t*_(*x*) in the landscape without making restrictive assumptions on its topography. This is necessary for the predictive analysis of real fitness landscapes, which do not have the symmetries of soluble models.

Note that (1) assumes that mutations occur independently of replication events. The results in this paper do not depend on this assumption: we could equally well consider Eigen’s quasispecies model [40], where mutations only arise as replication errors, or more general models (Methods). On the other hand, the infinite population limit implicit in (1) is a real limitation which overlooks the stochasticity of evolutionary histories. The applicability of deterministic models has been discussed extensively in the literature [40, 46], including from an experimental perspective [47]. Generally speaking, the infinite population approximation is good if the type space is low-dimensional or if the population is known to be localized in a small region of an otherwise high-dimensional genotypic landscape [48]. My own view is that deterministic models are a stepping stone towards any quantitative theory of evolutionary dynamics—we must understand how selection and mutation interact before we can ask about the influence of other evolutionary forces such as genetic drift.

### Classical analytical approaches

The first step to study (1) is to linearize it. This is commonly done by simply dropping the mean fitness term ⟨*ϕ*⟩_*t*_, the probability distribution *p*_*t*_(*x*) being then obtained from the solution *f*_*t*_ of

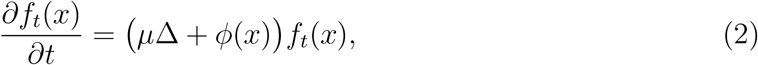

through normalization, *i.e.* 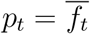 with 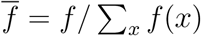. The linear equation (2) can then be solved formally in one of two classical ways—neither of which turns out to be directly useful for the prediction problem.

The first common approach to (2) uses the Feynman-Kac formula to write *f*_*t*_(*x*) as a weighted sum over Brownian paths *X*_*t*_ [49]. Unfortunately, these paths cover the whole fitness landscape, *i.e.* they are not by themselves predictive. Alternatively, we can decompose *f*_*t*_(*x*) over a basis of normal modes of the operator *A* = *μ*Δ + *ϕ* and consider the evolution of each component independently, as proposed by Eigen and Schuster [40]. This reduces (2) to a set of uncoupled growth equations, with the eigenvalues of *A* as growth rates. Accordingly, evolution is seemingly reduced to the natural selection of clouds of genetically related mutants, or “clans”.

The problem with the spectral approach is that, of all the modes of *A*, only one is positive and can therefore be interpreted as a distribution—the “quasispecies” distribution *Q*, i.e. the eigenfunction of *A* with the largest eigenvalue Λ. As a consequence, the Eigen-Schuster spectral approach is useful to characterize the asymptotic selection-mutation equilibrium *Q* = lim_*t→∞*_ *p*_*t*_, and in particular to determine whether this equilibrium is localized (adaptive) or delocalized (error catastrophe), but it cannot help us understand the approach to that equilibrium.

### Effective potential landscape

The key observation of this paper is that knowing *Q*—a single eigenfunction of *A*—to a good accuracy in fact goes a long way toward understanding evolutionary dynamics *far* from selection-mutation equilibrium. This is because from *Q* we can perform a change of variable that dramatically simplifies the analysis of evolutionary dynamics, as follows. Consider the function *g*_*t*_(*x*) = *e*^−Λ*t*^*Q*(*x*)*f*_*t*_(*x*), from which we can reconstruct the original distribution via 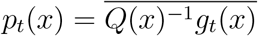. This function evolves according to

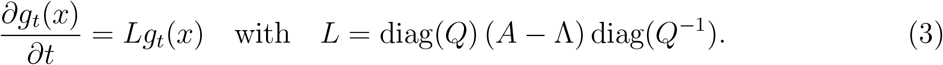

In Methods I show that, for any operator *A* that preserves the positivity of distributions, (3) is the forward Kolmogorov equation of a reversible *Markov process* with effective potential

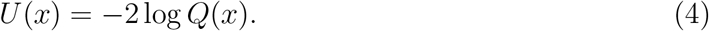

In the case where Δ is the Laplacian operator this process is just a biased random walk/Brownian motion:

- For discrete types, *L* generates nearest-neighbor jumps with transition rate

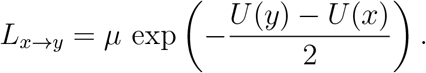
- For continuous types, *L* is the Fokker-Planck operator for a diffusion in the potential *U*, i.e.

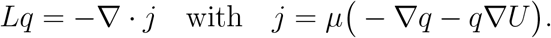

Note that the interpretation of the derived Markov process departs from that of the original selection-mutation model in two ways. First, *Q* is no longer viewed as coding the asymptotic equilibrium between selection and mutation, in which all transients are washed out; instead, (two times minus) its logarithm acts a potential landscape, whose role is to prescribe the dynamics *away* from equilibrium. Second, we are used to thinking of mutations as adding a random component to the otherwise deterministic flow of natural selection, with *μ* controlling the strength of the noise. Here, by contrast, *μ* plays the role of (*i*) an (inverse) time scale, and (*ii*) a parameter of the effective potential *U* which directs the evolution of the density in the space of types *X*. The noise component of the process itself has *unit* diffusivity.

What is the benefit of replacing the selection-mutation operator *A* by the Markov generator *L*? The answer is that the latter has an inbuilt notion of dominant evolutionary trajectory: from a given type *x*, the preferred path is the line of steepest descent of the effective potential *U*. Moreover, thanks to the smoothing effect of mutations imprinted in the quasispecies distribution, the potential landscape is far simpler—in particular, less rugged—than the fitness landscape itself. We now illustrate these aspects in more detail.

### Bare vs. effective ruggedness

As already mentioned, a classic approach to the ruggedness of fitness landscapes consists in counting the number of local fitness maxima [13]. For instance, in *NK* landscapes the expected density of fitness peaks grows from 2^−*N*^ (additive or “Mount Fuji” landscape) to (*N* + 1)^−1^ (uncorrelated or “house of cards” landscape) as the epistasis parameter *K* increases from 0 to *N* − 1, irrespective of the distribution of fitness components. However, the number of fitness peaks—the *bare* ruggedness of the landscape—is not directly relevant for evolutionary trajectories: at finite mutation rates, a low peak can be indistinguishable from no peak.

The results in the previous section imply that, in the large population limit, the evolutionary attractors are the local maxima of *Q* (local minima of *U*), not those of *ϕ*. But for a type *x* to be a local maximum of *Q*, it is not enough that its fitness be greater than that of its one-step mutants: an approximate expression for *Q* (given in (7) below and valid at low *μ*) shows that *ϕ*(*x*) must instead be greater than max *ϕ − μ*. This condition is typically much more stringent than the condition for *x* to be a local fitness maximum; the effective potential landscape is therefore significantly smoother than the fitness landscape. Thus, the number of *Q*-maxima of an NK lansdcape does not actually increase with *K*, but does with the skewness of the distribution of fitness components (data not shown).

### Reduced evolutionary dynamics

Next, the Markovian reformulation immediately suggests a coarse-grained (reduced) representation of evolutionary dynamics, as follows. For each local minimum *x_α_* of *U* we can consider the set of types *X_α_* from which *x_α_* can be reached along a *U*-decreasing path, its basin of attraction. The potential barrier between two adjacent basins is then given by *B_α→β_* = min_*π*_ max_*x*∈*π*_[*U*(*x*) − *U*(*x*_*α*_)] where *π* spans the directed paths connecting *X_α_* to *X_β_*. According to the standard Arrhenius-Kramers law for the transition time between minima of a potential landscape [43], the basin *X_α_* with frequency 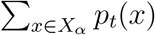 is *metastable* if

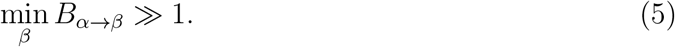

Large deviation theory further indicates that, of all the possible escapes from *X_α_* to an adjacent basin, the transition to argmin_*β*_ *B*_*α*→*β*_ is exponentially more likely to happen. This reduction in dynamical complexity is the main result of this paper.

The coarse-grained dynamics can be represented using tools usually applied to energy landscapes, such as the basin hopping graphs (BHG) recently developed in the context of RNA folding [50]. In a nutshell, a BHG is obtained by collapsing the local minima *x_α_* and their basins of attraction *X_α_* into nodes and connecting them according to adjacency relations between basins, weighted by the barrier height *B*_*α*→*β*_, as in Fig. 2.

### An evolutionary Lyapunov function

Finally, the Markovian reformulation provides a novel Lyapunov function for selection-mutation dynamics. An evolutionary Lyapunov function (ELF) traditionally refers to one of two distinct concepts. The first notion of ELF is a monotonic functional of distributions over type space *X*; examples include Fisher’s variance functional in the pure selection regime [51] or for type-independent mutation rates [52], or Sella and Hirsh’s free fitness functional in the SSWM regime [53] (see also [54]). The second kind of ELF is a monotonic functional of distributions over distributions over type space *X* (*i.e.* over allele frequency distributions); Iwasa’s [55] and Mustonen and Lässig’s [56] free fitness functions are of this kind.

Here I introduced a Markovian version of evolutionary dynamics in type space which is not restricted to pure selection or SSWM regimes. Since this Markov processes is reversible, the relative entropy (or Kullback-Leibler divergence) *D*[· || ·] with respect to its equilibrium distribution *α e*^−*U*^ = *Q*^2^ must decreases monotonically in time. This means that

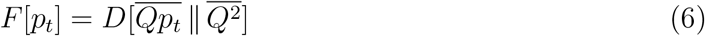

is a Lyapunov function for the evolutionary equation (1) for any mutation operator Δ and any mutation rate *μ* (Fig. 3). The construction of this ELF follows the same pattern as Iwasa’s and Mustonen and Lässig’s (as a relative entropy), but, unlike theirs but like Fisher’s, results in a functional of distributions over *X* and not allele frequency space. Also note that *F* [*p*_*t*_] is not merely an additive correction to mean fitness and thus goes beyond the scope of “free fitness” functions.

## EXAMPLES

To illustrate the predictive value of the Markovian formulation of selection-mutation dynamics we now consider two simulated fitness landscapes, chosen such that evolutionary attractors are not easily read off the landscape itself. (See Methods for explicit definitions.)

### Two-dimensional lattice

We begin with a two-dimensional rugged “phenotypic”[57] landscape, generated by sampling values from a Gaussian process with unit correlation length on a 30 × 30 lattice (with periodic boundary conditions). In the realization shown in Fig. 1A, the fitness landscape has a unique global maximum (green dot); this type corresponds to the maximum of the quasispecies *Q* for *μ* ≤ 0.02 (survival of the fittest), but not for higher mutation rates (survival of the flattest), see Fig. 1B.

**Figure 1.**
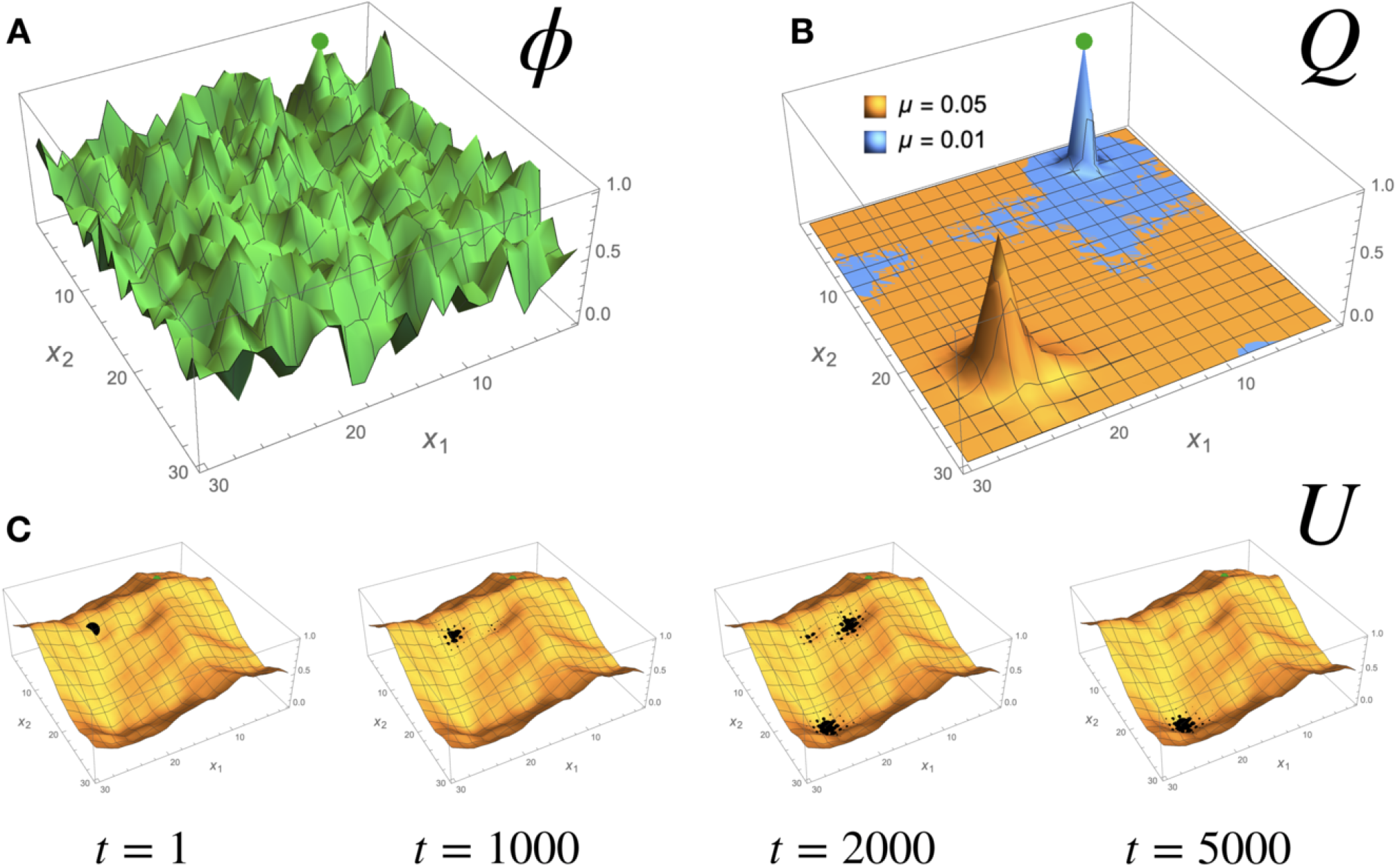
Evolution in a rugged 2d fitness landscape. A: The fitness landscape, obtained by sampling a Gaussian process with unit standard deviation and unit correlation length; the global fitness maximum is indicated by the green dot. It is *a priori* difficult to predict the path taken by a population evolving in this landscape. B: The quasispecies distributions *Q* for two different values of the mutation rate *μ*, localized at the fitness peak (low *μ*) or in some lower but flatter region (high *μ*). C: The effective potential *U* = −2 log *Q* for *μ* = 0.05 is much smoother than the fitness landscape, with few local minima which act as local attractors for an evolving population (black dots). Note how the population conspicuously moves away from the global fitness maximum.

Predicting the evolution of an initially monomorphic population directly from the topography of *ϕ* is clearly a difficult proposition. By contrast, examination of the effective potential *U* = −2 log *Q* (Fig. 1C) immediately reveals the preferred directions for its evolution: the population will go downhill in the potential *U*, potentially getting transiently trapped in the basins of its local minima and making transitions to other basins along the lowest saddles separating them. This is indeed the behavior of numerical solutions of the Crow-Kimura equation (Fig. 1C).

### Binary sequences with neutrality

As a simple model of a genotypic landscape with both ruggedness and neutrality, I considered an *NKp* landscape [58] of binary sequences with length *N* = 8, epistasis parameter *K* = 6 and neutrality parameter *p* = 0.7 (details in Methods). The landscape in Fig. 2 has 20 local maxima and an error threshold at *μ*_*c*_ ≃ 0.2. Comparing the basin hopping graphs of the fitness landscape *ϕ* and of the potential landscape *U* reveals that most of the complexity of the former is spurious. Moreover, coarse-grained evolutionary trajectories, described by the basin frequencies *p*(*X_α_*), is consistent with the succession of transitions predicated by the basin hopping graph of *U* : a population initially concentrated around the genotype 110 (a global fitness maximum) will evolve towards the flatter genotype 179 via the basins of 222 and 95 (Fig. 3A).

**Figure 2.**
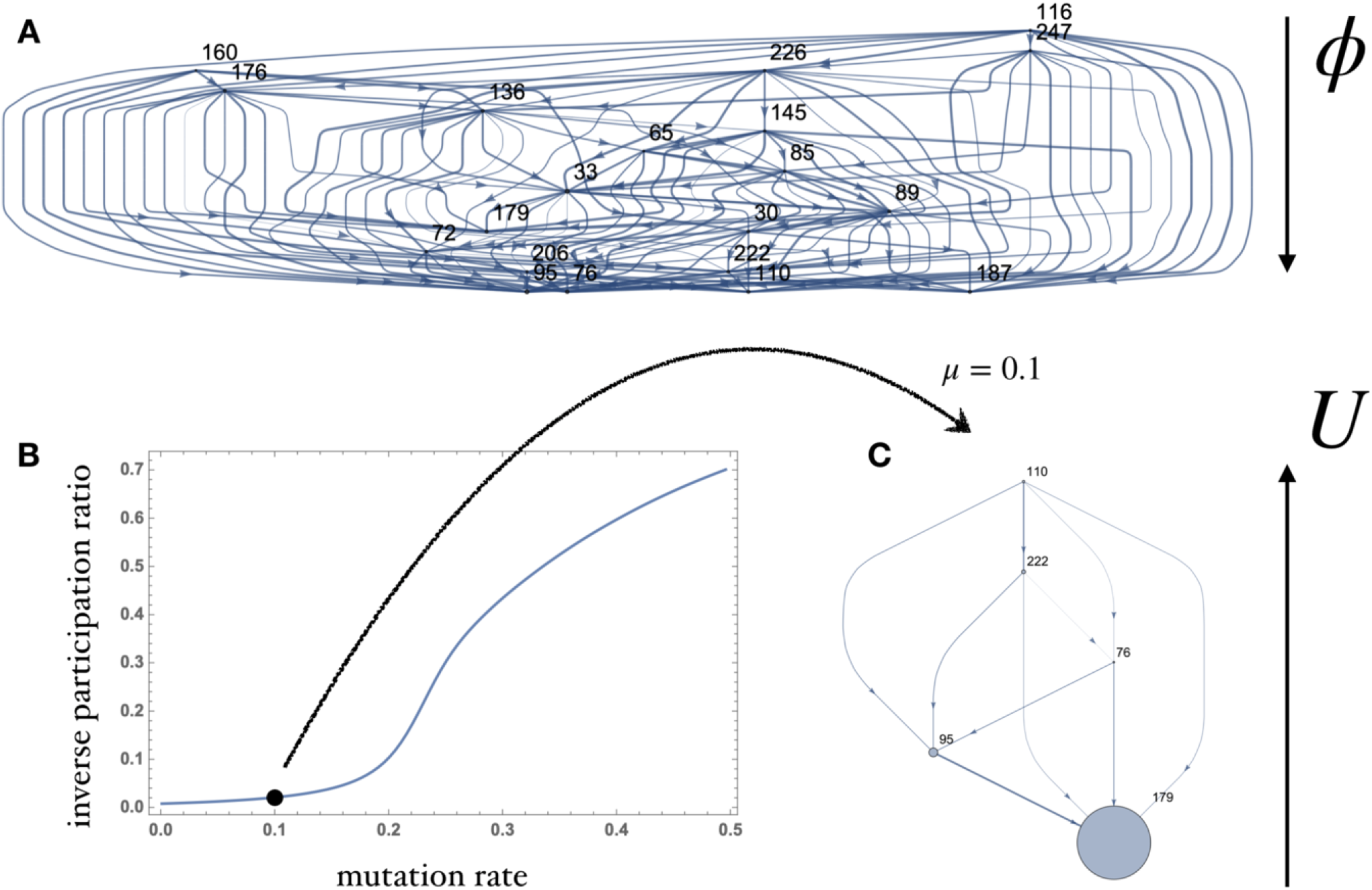
Evolution in an *NKp* genotypic landscape with 2^8^ = 256 types. A: The fitness landscape has 20 local fitness maxima and many saddles between them, making visualization and evolutionary prediction challenging. Here the landscape is represented as a basin hopping graph (BHG), in which nodes are basins of attractions of fitness maxima and edges adjacency relations between basins weighted by the barrier height. B: As the mutation rate passes a threshold at *μ* ≃ 0.2 (in units of the maximal fitness difference), the quasispecies distribution delocalizes, as signalled by the inverse participation ratio (∑_*x*_ *Q*(*x*)^2^)^−1^/|*X*|. C: The BHG for the effective potential (here for *μ* = 0.1) is much simpler—and immediately predictive, see Fig. 3.

**Figure 3.**
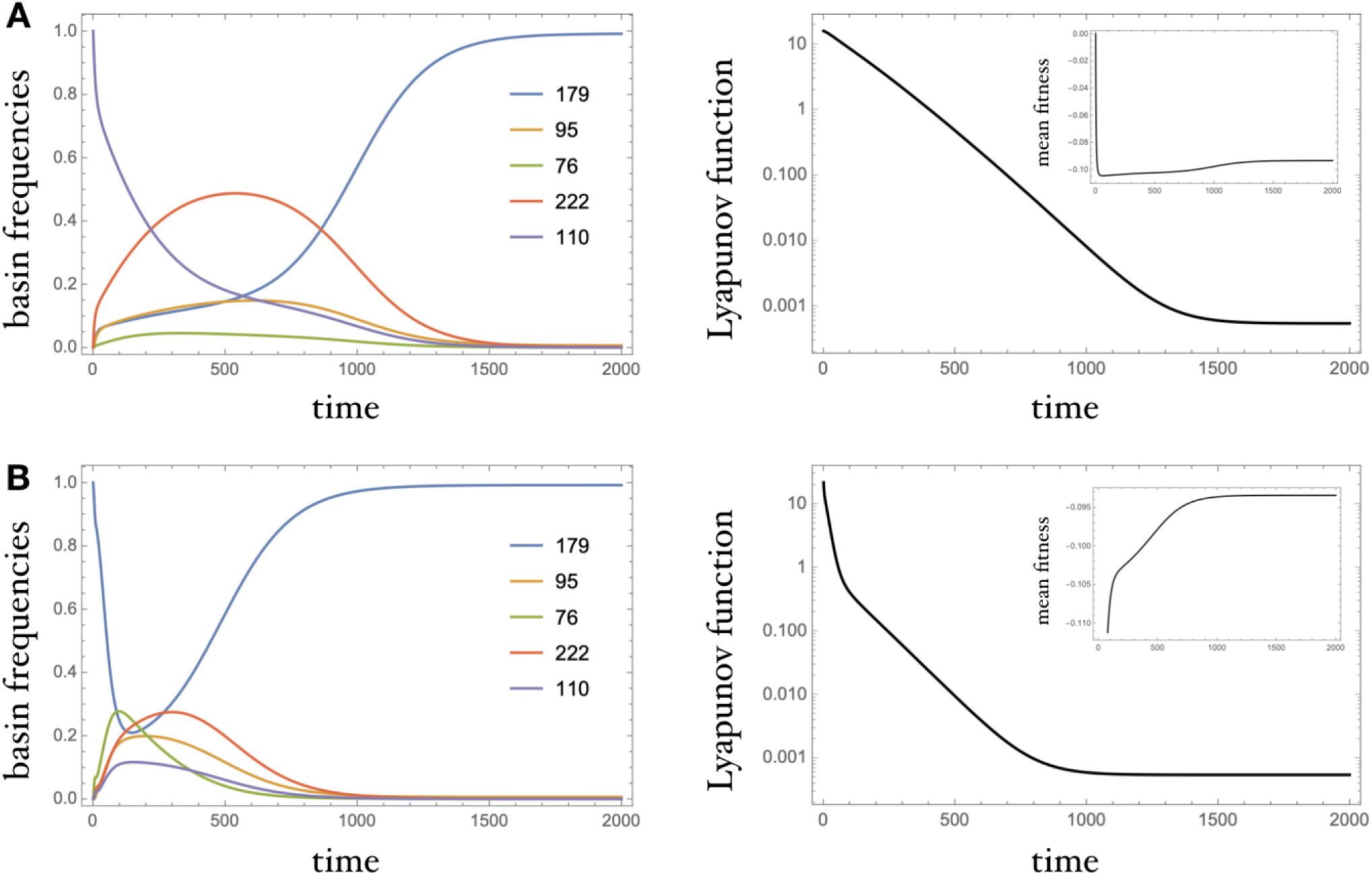
Evolutionary trajectories in the landscape of Fig. 2A, obtained by integration of the Crow-Kimura equation. A: A population initially concentrated in basin 110 moves towards basin 179 through basins 222 and 95, as suggested by the BHG in Fig. 2C. This happens in spite of the fact that 110 is a global fitness maximum and mean fitness decreases in time. B: Here the population starts off concentrated at type 179 and spreads in other basins under the effect of mutations, before returning to the basin of 179 as *t* → *∞*. This non-monotonic behavior of the basin frequency does not prevent the evolutionary Lyapunov function to decrease monotonically.

One also checks that the Lyapunov function (6) decreases monotonically also when the mean fitness ⟨*ϕ*⟩_*t*_ does not (Fig. 3A) and when the basin frequencies have strongly non-monotonic behavior (Fig. 3B).

## PHYSICAL ANALOGIES

Evolutionary theory has long benefited from analogies with statistical physics—the other field of science dealing with large, evolving populations—, see *e.g.* [53, 56, 59, 60]. More recently, Leuthäusser [61] and others [62, 63] have highlighted a parallel between evolutionary models in genotype space and certain *quantum* spin systems, which can be leveraged to compute the quasispecies distribution *Q* for some special fitness landscapes [45]. The present work was inspired by the realization that, in quantum mechanics, it *is* possible to map Schrödinger operators to diffusive trajectories—this is the basic idea underlying Nelson’s “stochastic” formulation of quantum mechanics [64].

But the scope of the analogy between evolution and non-equilibrium physics is, in fact, much broader: the interplay between selection and mutation is typical of *localization phenomena in disordered systems [65]*, be them classical or quantum. The linearized Crow-Kimura equation 2, for instance, is formally identical to the parabolic Anderson model [49, 66, 67], a simple model of intermittency in random fluid flows; the linearized Eigen model in turn resembles the Bouchaud trap model [68], a classical model of slow dynamics and ageing in glassy systems. These physical phenomena have obvious evolutionary counterparts: the Anderson localization transition corresponds to the error threshold; intermittency to epochal or punctuated evolution; tunnelling instantons to fitness valley crossings; and ageing to diminishing-return epistasis. The generalization of Nelson’s mapping of the Scrödinger equation to a diffusion process presented in this paper implies that all are in fact unified under the familiar umbrella of Markovian metastability.

The value of such analogies is twofold. First, they bring the large repertoire of results and techniques derived in condensed matter and nonequilibrium physics to bear on evolutionary dynamics. As an elementary example, we can use the forward scattering approximation (FWA) familiar from Anderson and many-body localization theory [69] to compute the effective potential *U* in the small *μ* limit, as the (log of the) ground state of the quantum Hamiltonian *H* = −*A* = −*μ*Δ − *ϕ*; for non-degenerate fitness landscapes this gives

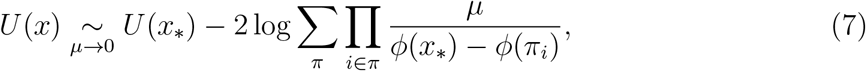

where *x*_∗_ = argmax *ϕ* and *π* spans the set of shortest paths between *x*_∗_ and *x*. This gives suprisingly good results, including for large mutation rates (Fig. 4).[70] Conversely, the link between evolution and the physics of disordered media can stimulate new work in physics and mathematics. As already mentioned, the generator of selection-diffusion dynamics is not always Hermitian (it is not in Eigen’s model). This suggests that some of the results usually derived for random Schrödinger operators can likely be generalized for more general classes classes of operators, as already emphasized by Altenberg [71].

**Figure 4.**
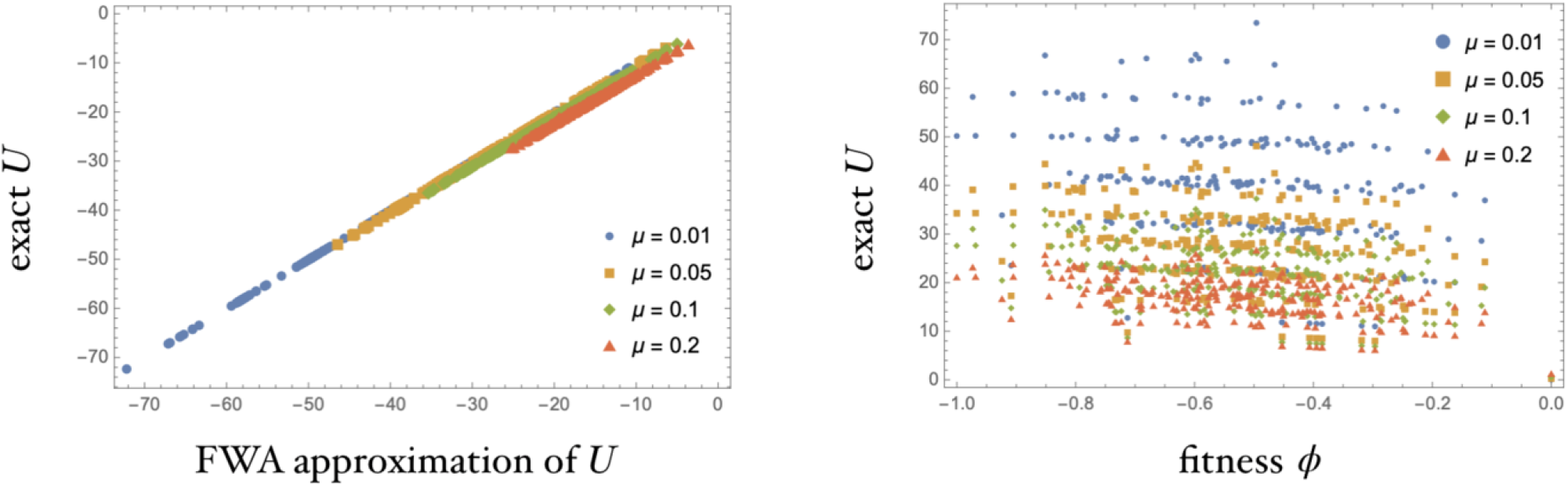
Effective potential for a non-degenerate NK landscape with *N* = 8 and *K* = 6. The FWA approximation familiar from Anderson localization theory gives excellent results, including at large mutation rates (left). By contrast, the bare fitness values *ϕ* are poorly correlated with the effective potential *U* (right). Here mutation rates are given in units such that *ϕ* ranges from−1 to 0. -

## CONCLUSION

A widely shared understanding of the role of mutations in evolution has them feeding raw material to the fitness-maximizing sieve of natural selection. But when mutation rates are high, as they are in *e.g.* RNA viruses [72] and likely were in early life [40], evolutionary success requires more than the discovery of a high-fitness mutant genotype: the mutants of the new mutant must also have relatively high fitness, *i.e.* the mutant type must be mutationally robust. The effective potential *U* introduced in this paper combines fitness and flatness into a single evolutionary potential—should we call it “flitness”?—which directly determines evolutionary trajectories across the spectrum of mutation rates. I argue that instead of the fitness landscape itself, it is this effective potential that we should analyze, coarse-grain, etc. if we are to predict evolution.

On a conceptual level, the effective potential *U* addresses two longstanding questions in evolution: *(i)* On what time scale (individual generation, infinite lineage) should “fitness” be defined [73]? and *(ii)* What quantity does evolution optimize [74]? My proposed answers are, respectively: *(i)* It is fine to define the fitness *ϕ* (*g*) of a type *g* as reproductive success over one generation, which makes it directly measurable, but one should keep in mind that *ϕ* (*g*) is not necessarily a good predictor for the success of a lineage descending from *g*—this role is played by the effective potential *U* (*g*); and *(ii)* like other dissipative processes, evolution through selection and mutations minimizes the statistical divergence to its Markovian equilibrium. There is an arrow of time in micro-evolution—just not one that points towards maximal fitness.

## ACKNOWLEDGEMENTS

This work was inspired by the results of Filoche and Mayboroda on wave localization [75] and those of Yasue on quantum tunnelling [76]. I am indebted to M. Kenmoe, A. Klimek, M. Lässig, O. Rivoire and D. Saakian for discussions and feedback. Funding for this work was provided by the Alexander von Humboldt Foundation in the framework of the Sofja Kovalevskaja Award endowed by the German Federal Ministry of Education and Research.

## METHODS

### From positive to Markov semigroups

The main result of this paper is best formulated in terms of positive operator semigroups [77]. A positive operator semigroup (*P*_*t*_)_*t*≥0_ is one that preserves the positivity of distributions on a space *X*, but not their normalization. This is the case of the linear flow (*P*_*t*_) = (*e*^*At*^) if the non-diagonal elements of its generator *A* are all non-negative (i.e. *A* is “essentially non-negative”). Up to the addition of a multiple of the identity, we may further assume that the diagonal elements are also non-negative, i.e. *A* is a non-negative operator.

The Perron-Frobenius theorem states that *A* has a left eigenvector *Q* with simple eigen-value Λ whose components are all positive in each irreducible component; moreover *P*_*t*_ = *e*^*At*^ converges to the projection operator on *Q* as *t* → *∞*. Now, under the conditions above, the operator

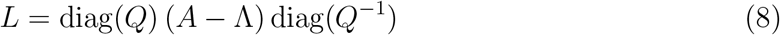

is the infinitesimal generator of a reversible Markov process on *X* with equilibrium distribution ∝ *e*^−*U*^ with *U* = −2 log *Q*. This is easily proved as follows.

If *X* is a discrete space (genotypic landscape), we must check that *L* satisfies the conditions for a transition rates matrix, namely that *L* has non-negative off-diagonal elements and ∑_*i*_ *L*_*ij*_ = 0. The former follows from the same property for *A* because 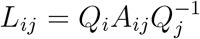 for *i* ≠ *j*. The latter follows from *Q* being a left eigenvector of *A* with eigenvalue Λ:

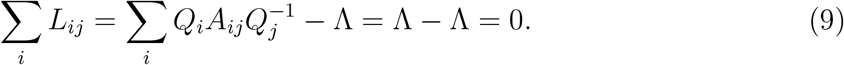

Note that, when *A* = *μ*Δ + *ϕ* with Δ the Laplacian on a graph (such that Δ_*ij*_ = 1 when *i* and *j* are adjacent and zero if *d*(*i, j*) > 1), then *L* generates nearest-neighbor jumps with rate 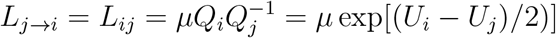, as stated in the main text.

For the continuous case, consider a domain of ℝ^*d*^ and assume for simplicity that the mutation operator Δ = ∇^2^ is the Laplacian in that domain, generating a standard *d*-dimensional Brownian motion. In this way *A* is a self-adjoint Schrödinger operator. Let. 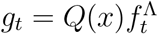, where 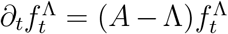. An explicit computation then shows that *g*_*t*_ satisfies the continuity equation *∂_t_g_t_* = −∇.*j*_*t*_ with the reversible flux *j*_*t*_ = −*μ*(∇*g*_*t*_ − *g*_*t*_∇*U*). This is the Fokker-Planck equation for a diffusion process with unit diffusivity and potential *U*.

### Model landscapes

The Gaussian process landscape of Fig. 1 is obtained by sampling a vector from the multivariate Gaussian distribution with zero mean and *L*^2^ × *L*^2^ covariance matrix *G*_*x,y*_ = *e*^−*d*(*x,y*)^ where *d* denotes the distance function on the two-dimensional periodic lattice ℤ_*L*_ × ℤ_*L*_.

The *NK_p_* fitness landscape over the hypercube {0, 1}^*N*^ with epistasis (or ruggedness) parameter *K*, neutrality parameter *p* and component distribution 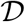 is defined by the formula 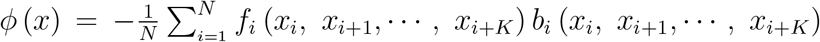 where the components of the binary string *x* are identified cyclically and the values of functions *f*_*i*_, *b*_*i*_ : {0, 1}^*K*+1^ → ℝ are i.i.d. samples from 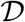 and Bernoulli(1 − *p*), respectively. Unless specified otherwise it is customary to take 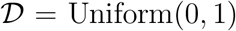. The *NK* model is the special case when *p* = 0, *i.e.* without neutrality.

